# The effect of genome organisation on selection efficiency in two contrasted plant species

**DOI:** 10.64898/2025.12.19.695387

**Authors:** Jennifer E. James, Martin Lascoux

## Abstract

Does the distribution of fitness effects of new mutations vary across the genome? Under the classical Fisher Geometric Model (FGM) we might not expect it to. In FGM, phenotypic traits are envisioned as dimensions of a landscape, with fitness determined by position in the landscape, i.e., the particular combination of traits of an individual. New mutations are represented by vectors that move from an ancestral to a new phenotype. In classical FGM these vectors affect all trait dimensions simultaneously (’universal pleiotropy’). However, introducing partial and modular pleiotropy into an FGM framework leads to an expectation that parameters of the DFE will vary with mutational pleiotropy-the number of traits affected by individual mutations. Here we address this prediction by investigating whether traits related to mutational pleiotropy, expression level and network connectivity, affect the parameters of the DFE using whole genome data from A. thaliana and C. grandiflora, two closely related Brassica species that vary significantly in their demography and mating system, and therefore, in effective population size and the effects of linked selection. Results were similar across both species. We found that expression level and network connectivity were predictive of the parameters of the deleterious DFE, even once co-correlations among genome biology traits were accounted for. Our results suggest that, across the genome, molecular evolutio(high mutational pleiotropy). nary patterns agree with the predictions of FGM, albeit relaxing the assumption of universal pleiotropy, and that variation in mutational pleiotropy among genes is sufficient to have detectible effects on the DFE.

**Significance statement:** How do the effects of new mutations vary across the genome? If mutations in some genes affect many traits (high mutational pleiotropy), we hypothesise they will be more strongly deleterious, with lower variance in their selective effects. We test this by investigating the distribution of effects of new mutations across genes that vary in features that are related to mutational pleiotropy: expression level, gene network connectivity, and number of associated GO terms. The mean strength and coefficient of variation of selection of new mutations varied across genes with different features in the manner expected by our hypothesis. This demonstrates that important parameters of molecular evolution can vary across the genome with genome architecture.

## Introduction

The Fisher Geometric Model (FGM) (Fisher, 1930) is a widely used framework that has proved extremely useful for understanding the effects of mutations. Under the FGM, the traits of an organism are imagined as independent dimensions of a fitness landscape; an individual with a given phenotype can therefore be represented by a point in this landscape. We assume there is an ‘optimum’ point in the fitness landscape defined by a particular combination of trait values, i.e., individuals close to the optima are more fit than those further away. Mutations can then be envisioned as vectors between the ancestral and the new phenotype, with mutation strength represented by vector length. Mutations in this model affect all trait dimensions and are thus described as exhibiting universal pleiotropy. Mutations that move individuals closer to the optimum are beneficial, those that move them further away, deleterious.

In classical FGM, the number of dimensions of the fitness landscape is the phenotypic complexity of the organism. Fisher himself used the FGM to demonstrate that the mean strength of selection acting on new deleterious mutations will increase with increasing complexity, because with increasing dimensionality, it is more likely that a new mutation will move an organism away from the fitness optima in at least some dimensions of the phenotypic space. Put simply, a new mutation is more likely to disrupt a complex system than a simple one. Orr (2000) extended this analysis to show that as the number of phenotypic traits increases, there is also a reduction in the speed of adaptation, a phenomenon known as the ‘cost of complexity’.

Therefore, the FGM provides one hypothesis for why we expect the average strength of mutations to vary with varying levels of complexity. However, one of the key assumptions of FGM, universal pleiotropy, may not hold, and the extent of pleiotropy across biological systems is generally a matter of some debate (see, for example, (Hill & Zhang, 2012a, 2012b; Wagner et al., 2008; Wagner & Zhang, 2011)). While pleiotropy appears to be pervasive, the genetic architecture of traits may be modular, such that certain genes interact more highly with each other, and are more closely associated with particular phenotypic traits, than others (Mackay & Anholt, 2024). The extent to which genes are organised into these functional modules that produce traits, and how much genes interact across modules, is a topic of ongoing research (Aguirre et al., 2025b, 2025a).

Introducing partial and modular pleiotropy into the FGM framework has interesting consequences for our predictions of patterns of molecular evolution across the genome (Chevin et al., 2010; Lourenço et al., 2011). In such models, pleiotropy is defined as mutational pleiotropy: the number of traits affected by a mutation. Restricted pleiotropy was found to strongly affect the expected distribution of fitness effects of new mutations (the DFE). For populations at the fitness optimum, the mean effect of mutations scales with the pleiotropy of mutations, not phenotypic complexity, such that if there is high pleiotropy, mutations have stronger average effects and the variation in the fitness effects of new mutations is low. To put this in terms of traits: if mutational pleiotropy is high, every new mutation that affects fitness will affect every trait, but as mutational pleiotropy decreases, new mutations will affect lower numbers of traits, therefore, the fitness effects that new mutations have will be lower on average and have higher variability in their effects as only some traits are affected. If these models are correct, then we expect the DFE to vary across the genome with the level of pleiotropy.

There is evidence that the DFE does vary across the genome with features of genome biology. For example, it has long been known that one of the strongest predictors of the rate of protein evolution is expression level, such that highly expressed genes evolve slowly, while other predictors, such as proposed functional importance, appear to have a weaker effect on evolutionary rate (Pál et al., 2001; Wright et al., 2004; Zhang & He, 2005; Zhang & Yang, 2015). This has been interpreted as evidence that more highly expressed genes experience greater selective constraint due to a greater risk of mistranslation and subsequent protein misfolding, which can be cytotoxic to cells (Drummond & Wilke, 2008), not necessarily because there is greater purifying selection due to greater functional constraint on highly expressed genes. While this misfolding-avoidance hypothesis has been called into question, for example, by the finding that highly expressed proteins are not more thermodynamically stable (Plata & Vitkup, 2018), it has been supported by recent experimental work. Wu et al. (2022), who performed targeted mutagenesis experiments to investigate the fitness landscapes of mutations in two genes in yeast, found that a large proportion of the effects of new mutations were dependent on the gene’s expression level, with mutations that lead to protein misfolding being particularly deleterious. Although this work involved only two genes, the results highlight the importance of considering how genome biology traits moderate the effects of new mutations from a fitness landscape perspective.

Here, we explore the extent of within-genome variation in the parameters of the DFE across two closely related, yet very different, Brassica species: *A. thaliana*, a cosmopolitan selfing species with low genetic diversity (Igolkina et al., 2025); and *C. grandiflora*, an outcrossing species with a limited geographical range and very high genetic diversity (Williamson et al., 2014). The aim of this study is to provide a more general overview of the effects of genome biology on the fitness landscape of new mutations, to assess whether findings agree with previous results and accord with expectations from the FGM. We are therefore particularly interested in testing a possible relationship between the parameters of the DFE and gene properties that relate to pleiotropy. We thus focus on gene network connectivity, which is perhaps the most commonly used proxy of pleiotropy across the literature (see, for example, Josephs et al., 2017; Rennison & Peichel, 2022; Ruelens et al., 2023). We also consider the effect of expression level. More highly expressed genes have been found to have more pleiotropic effects (Barbitoff et al., 2025; Guillaume & Otto, 2012), and, as detailed above, expression level is known to affect the fitness landscape of new mutations.

## Results

### The effects of new mutations and proxies for mutational pleiotropy

We first considered how two proxies of mutational pleiotropy, expression level and gene network connectivity, affect the mean strength and variance of selection acting on new mutations. For this we inferred the distribution of fitness effects of new mutations (DFE). The DFE is estimated from the site frequency spectrum of neutral and non-neutral sites (here, four-fold and zero-fold synonymous sites) and is assumed to be a mixture between gamma (for deleterious mutations) and exponential (for beneficial mutations). The parameters that govern the deleterious gamma DFE are shape (*b*) and scale (*θ*); for interpretability we here report the mean strength of selection acting against deleterious mutations, *S_d_*, the shape parameter of the deleterious DFE, *b*, and the coefficient of variation in the mean strength of selection acting against new mutations, 1⁄√*b* (Tataru et al., 2017).

DFE inference presents a problem at the gene level: per gene, there are relatively few SNPs, resulting in stochastically noisy SFSs from which it would be difficult to infer the DFE reliably. To get around this problem, we therefore used a grouping strategy. For example, for expression level, we first ranked genes by their expression level. Ranked genes were then divided into groups of different expression levels, such that each group had approximately the same number of 4-fold synonymous polymorphisms (*p_4_*). This also resulted in each group containing similar numbers of genes. We here present results for genes in 20 groups, although results were confirmed for different numbers of groups (Suppl. Figs. 1 and 2).

**Figure 1).**
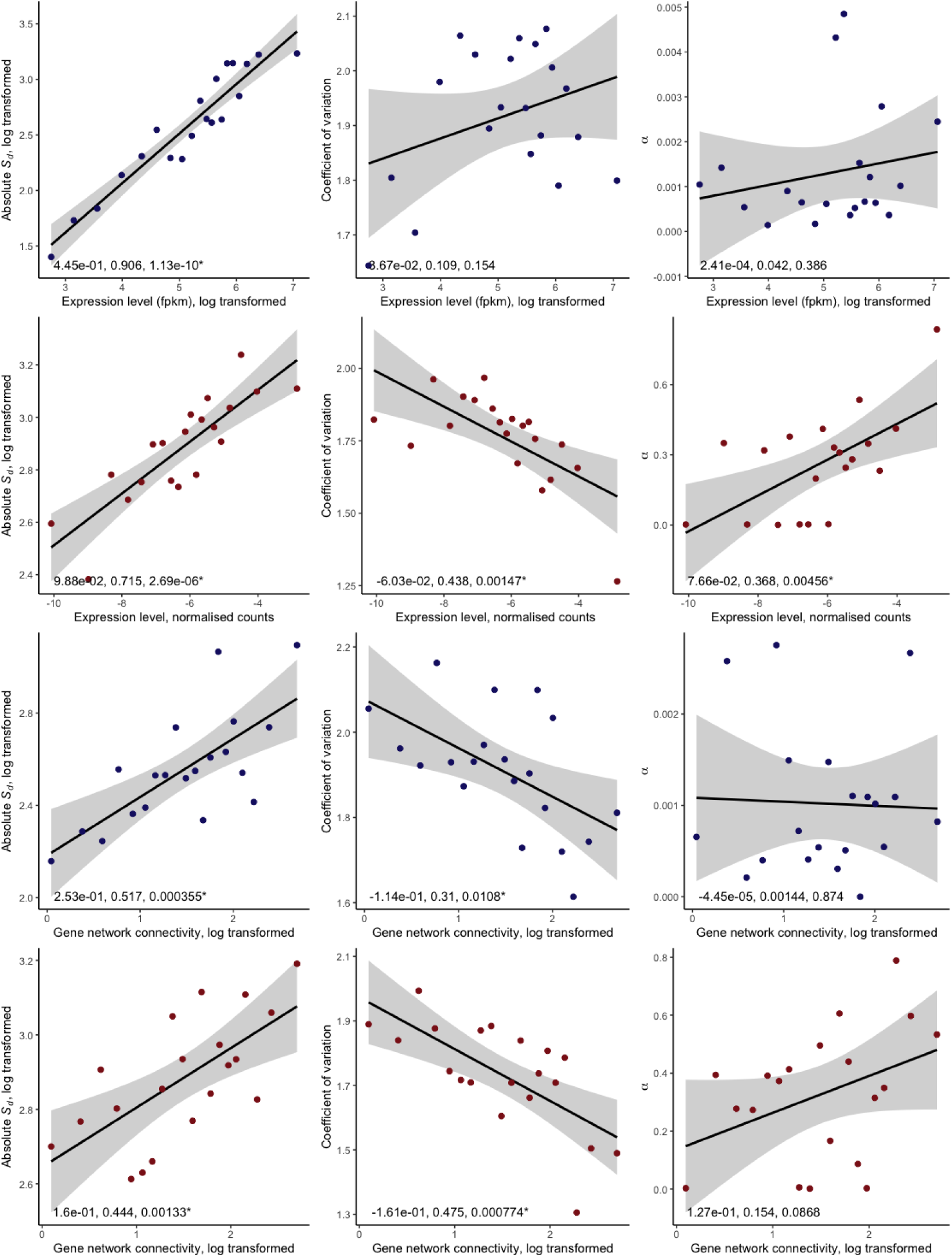
Parameters of the DFE and two genome traits related to pleiotropy; expression level and gene network connectivity. The first column shows the absolute mean strength of selection acting against new deleterious mutations, *S_d_* (the scale parameter of the DFE), the second shows the coefficient of variation of selection acting against new deleterious mutations (related to the shape parameter *b* of the DFE), the third shows *α*, the proportion of substitutions estimated to be fixed by adaptive evolution, as estimated from the beneficial DFE (i.e., using polymorphism data only). Rows indicate different genome traits: either expression level or the estimated level of connectedness in functional gene interaction networks, estimated from number of gene interaction partners. Colour indicates species; dark blue: *A. thaliana*, dark red: *C. grandiflora*. Points show the estimated value of the estimated DFE parameter, estimated over all genes in that bin, plotted against the mean value of the genome trait for that bin. Lines are linear regression results (log transformed genome biology traits), shown with 95% confidence intervals. Results of linear regression analyses are shown on each plot (slope, *R^2^*, and *p* value), with results significant at the p < 0.05 level marked ‘*’.

**Figure 2).**
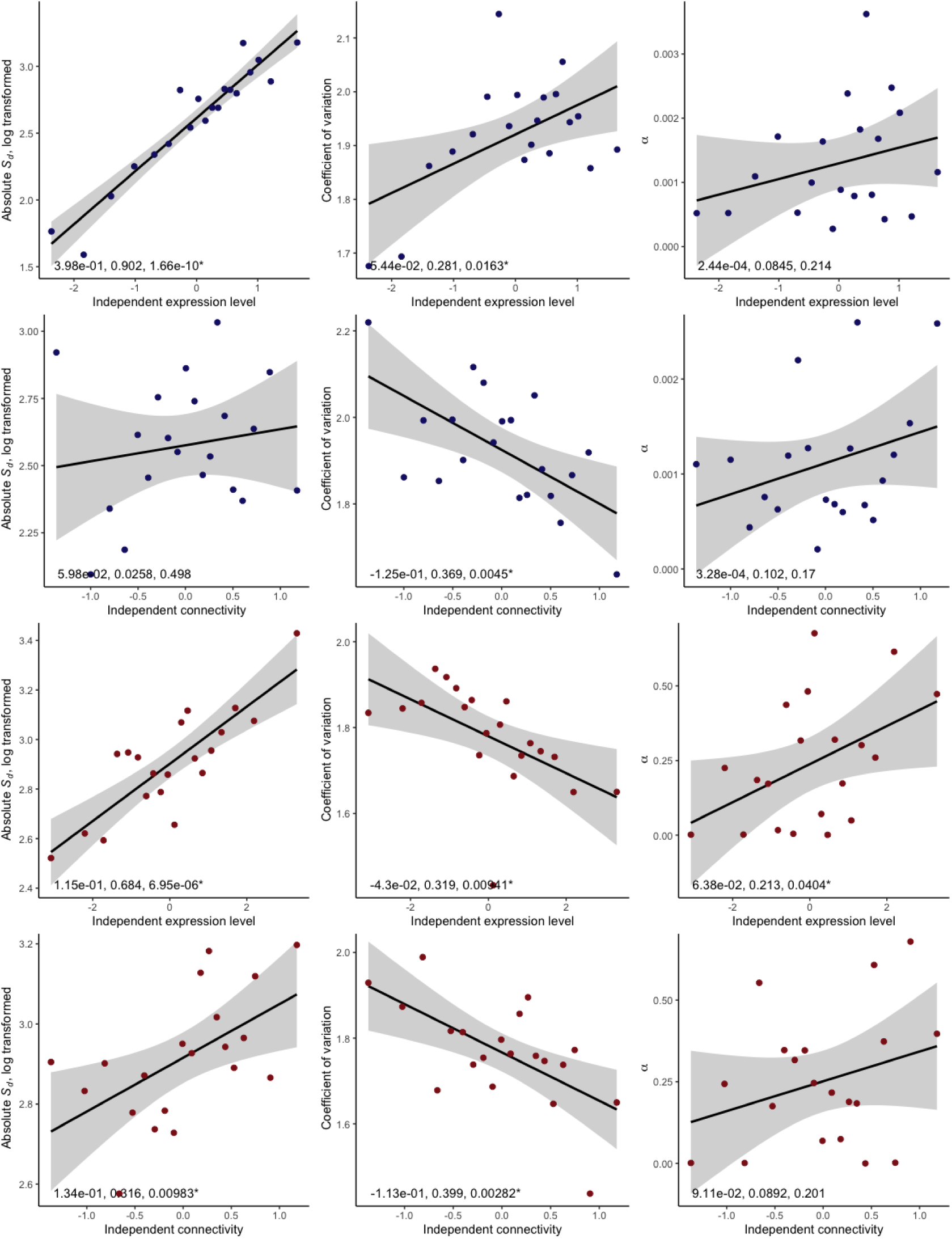
Parameters of the DFE and two genome traits related to pleiotropy, expression level and gene network connectivity, after correction for their association with other gene features, making them statistically independent. The first column shows the absolute mean strength of selection acting against new deleterious mutations, *S_d_* (the scale parameter of the DFE), the second shows the coefficient of variation of selection acting against new deleterious mutations (related to the shape parameter *b* of the DFE), the third shows *α*, the proportion of substitutions estimated to be fixed by adaptive evolution, as estimated from the beneficial DFE (i.e., using polymorphism data only). Rows indicate the genome traits after correction: either expression level or the estimated level of connectedness in functional gene interaction networks, estimated from number of gene interaction partners. Colour indicates species; dark blue: *A. thaliana*, dark red: *C. grandiflora*. Points show the estimated value of the estimated DFE parameter, estimated over all genes in that bin, plotted against the mean value of the corrected genome trait for that bin. Lines are linear regression results (log transformed genome biology traits), shown with 95% confidence intervals. Results of linear regression analyses are shown on each plot (slope, *R^2^*, and *p* value), with results significant at the p < 0.05 level marked ‘*’.

**Figure 3).**
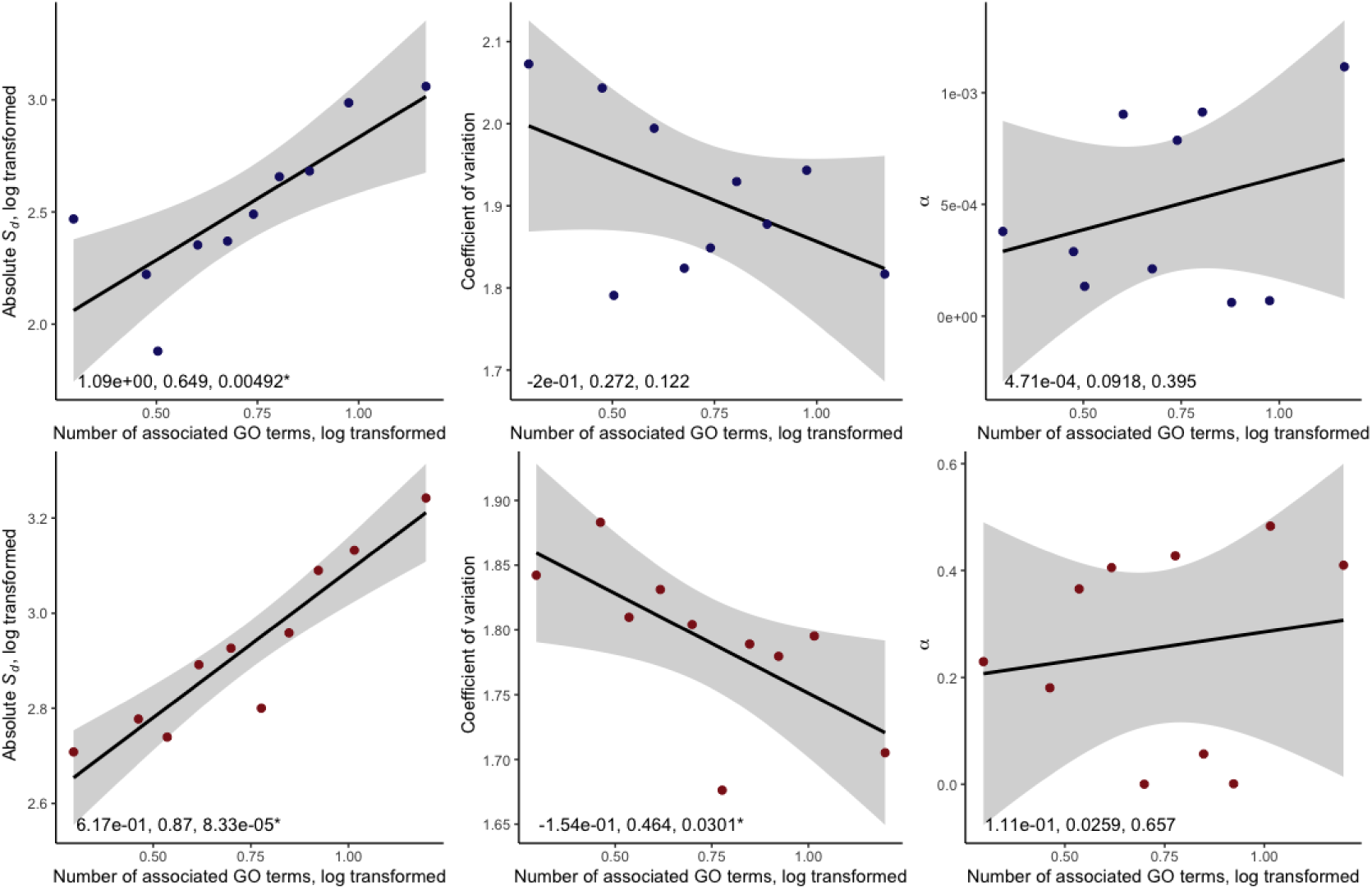
Parameters of the DFE and the number of GO terms associated with each gene. The first column shows the absolute mean strength of selection acting against new deleterious mutations, *S_d_* (the scale parameter of the DFE), the second shows the coefficient of variation of selection acting against new deleterious mutations (related to the shape parameter *b* of the DFE), the third shows *α*, the proportion of substitutions estimated to be fixed by adaptive evolution, as estimated from the beneficial DFE (i.e., using polymorphism data only). Colour indicates species; dark blue: *A. thaliana*, dark red: *C. grandiflora*. Points show the estimated value of the estimated DFE parameter, estimated over all genes in that bin, plotted against the mean number of associated GO terms for all genes in that bin. Lines are linear regression results (log transformed genome biology traits), shown with 95% confidence intervals. Results of linear regression analyses are shown on each plot (slope, *R^2^*, and *p* value), with results significant at the p < 0.05 level marked ‘*’.

We found that both expression level and gene network connectivity were predictive of the parameters of the DFE. Across both the *A. thaliana* and *C. grandiflora* genome, mutations in genes that had more functional network connections (as predicted by AraNet (Lee et al., 2015)) were on average more strongly deleterious, and the coefficient of variation in selection strength was lower. Similarly, we found that in *C. grandiflora*, mutations in genes that had higher expression levels had a higher mean deleterious selection coefficient and a lower coefficient of variation in selection strength. However, in *A. thaliana*, we found that expression level correlates positively with the mean deleterious selection coefficient acting on new mutations, but is not significantly related to the coefficient of variation.

Several gene features are known to be correlated (see Suppl. Fig. 3 for details of how the gene metrics are correlated). Co-correlations among variables might drive some of the relationship we see between expression level and gene network connectivity and the parameters of the DFE. We therefore conducted multiple linear regression models for both expression level and gene network connectivity with other gene features and took the residuals of these models, thereby generating metrics of expression level and gene network connectivity that are uncorrelated to other features. We then repeated our gene ranking and DFE inference steps. We found that for these independent measures of expression level and gene network connectivity, we recover very similar relationships with the mean strength and coefficient of variation of selection acting on new deleterious mutations.

While gene network connectivity and expression level are commonly used proxies for pleiotropy, a more direct estimate of pleiotropy would estimate the number of specific phenotypes affected by each gene. One way to infer this information is to investigate the number of different GO terms associated with each gene. Different GO terms represent different molecular functions and therefore different phenotypes; an association with more GO terms could therefore represent a more direct association between pleiotropy than expression level or gene network connectivity. There is relatively little variation in the number of associated GO terms across genes, making it difficult to perform our grouping strategy for high numbers of bins; however, for both *A. thaliana* and *C. grandiflora*, we found that an association with a higher number of GO terms resulted in stronger purifying selection acting on new deleterious mutations, and a lower coefficient of variation on the strength of selection acting on new deleterious mutations.

### Adaptive evolution

In *A. thaliana*, *α*, the estimated proportion of substitutions that are adaptive (also referred to as *α_DFE_*, estimated from polymorphism data only) was uniformly low across the dataset with a mean of 0.001, and we found little evidence that genome biology traits were predictive of *α*. However, in *C. grandiflora* there was considerably more evidence of adaptive evolution across the genome, with an estimated proportion of 12% of substitutions fixed through adaptive evolution, which is likely a consequence of the more efficient selection across the genome of this outcrossing plant species as compared to the selfing *A. thaliana*. While only one genome biology trait (expression level) is correlated with *α* at the 0.01 level, relationships with our likely proxies of pleiotropy are always positive in *C. grandiflora*. As *α* is the proportion of substitutions that are adaptive, this may reflect a lower proportion of non-adaptive substitutions being fixed due to more efficient purifying selection, rather than a higher rate of adaptation *per se*.

### Per gene metrics

To further explore these results conducted on bins of genes, we also considered patterns in the efficiency of selection with gene features at the per-gene level. We cannot calculate the DFE for individual genes due to insufficient polymorphism data; however, we are able to calculate *π_0_* and *π_4_* per gene, and investigate how *π_0_*/*π_4_*, a measure of the efficiency of selection, varies with genome biology features. In all the following, we report results from scaled genome biology features, while *π_4_* was Box-Cox transformed and *π_0_*/*π_4_* was log transformed, as appropriate to better approximate normally distributed residuals.

In our dataset, genome biology factors are highly correlated; we therefore considered large linear models that incorporated interaction terms among all fixed effects. However, for *A. thaliana π_4_* these did not significantly improve the fit of our model. We found that expression level, gene network connectivity, recombination rate and protein length are predictive of *π_4_*, with length and recombination rate being particularly important. Alone, these two variables explain 2.11% of the variance in *π_4_*, while the full model explains 2.37% of the variance.

In contrast, connectivity and GO term count, traits both associated with pleiotropy, are the strongest predictors of *π_0_*/*π_4_* across the *A. thaliana* genome, with our model including all gene features explaining 4.64% of the variance in *π_0_*/*π_4_* overall. Incorporating interaction terms into this model did improve the fit to the data, albeit resulting in only a small improvement in the amount of variance we explained (5.19% overall). The increase in the explanatory power of the model primarily had to do with the incorporation of only two interaction terms: the interaction between expression level and length (negative), and expression level and GO term count (positive), resulting in a model which explained 5.02% of the variance in *π_0_*/*π_4_*. For *π_0_*/*π_4_*, the fixed effects with the highest estimated contribution were expression level, GO term count, and connectivity. Removing any of these variables resulted in significantly worse model fit and a drop in the explanatory power of our model (3.74% of the variance in *π_0_*/*π_4_* was explained without connectivity, 4.27% without expression level, 3.23% without GO term count).

In *C. grandiflora*, we have fewer gene features, however, results are qualitatively similar. We find that of expression level, gene network connectivity, GO term count and protein length, only length is significantly predictive of *π_4_* (such that a model including connectivity, expression level and length is not a significantly better fit to the data than a model including just length, p = 0.32), explaining 4.11% of the variance. By contrast, all gene features are predictive of *π_0_*/*π_4_* in *C. grandiflora*; in combination they explain 9.69% of the variance. Incorporating interactions between the variables also significantly improved the model fit, but only the interaction between expression level and gene network connectivity (negative) was significant, resulting in a best model that explained a total of 9.78% of the variance in *π_0_*/*π_4_*. Again, removing any of the gene features resulted in a significantly worse model fit, with traits associated with pleiotropy having a stronger effect; without length, 9.26% of the variance in *π_0_*/*π_4_* was still explained by the model, while without GO term count this dropped to 8.81%, without connectivity, 8.18%, and without expression, 7.31%. Overall, our per-gene results agree with our DFE results from binning genes. Additionally, they suggest that a combination of gene property variables is better able to explain the variance in the efficiency of selection across the genome than any one gene property alone.

## Discussion

In this study we have investigated how genome biology affects patterns of molecular evolution, and whether traits that are proxies of pleiotropy impact the fitness effects of new mutations. We find that both expression level and gene network connectivity affect the mean and coefficient of variation of the strength of selection acting on new mutations, and that this effect holds when the correlation between these variables is corrected for. Our analysis demonstrates that the parameters of the DFE are likely to vary systematically across the genome with gene features, and that this variation is largely in agreement with our hypotheses based on the predictions of FGM.

We conducted our analysis on two plant species, *A. thaliana* and *C. grandiflora*. To have sufficient resolution across the genome to perform per-gene polymorphism analyses, we chose *A. thaliana* because it has a high number of sequenced individuals, in addition to being a model plant species with an extremely well-studied genome. *C. grandiflora*, as a close relative, has a similar genome structure and gene content. While there are fewer sequenced *C. grandiflora* individuals that we could include in this analysis, this species has a high degree of genetic diversity, which again allows us to estimate population genetics statistics on a per-gene basis. Unlike other model species and their close relatives, *A. thaliana* and *C. grandiflora* are very different in their demography and life history traits, making them an interesting pair of species to contrast. Selfing species such as *A. thaliana* are known to vary from their outcrossing relatives, like *C. grandiflora*, in effective population size and the effectiveness of selection (Charlesworth, 2003); additionally, reproductive mode has been shown to have an impact on the DFE in plants (Chen et al., 2020), and, as expected the two species have very different DFEs overall despite their phylogenetic relatedness. We also infer the proportion of adaptive substitutions to be much higher in *C. grandiflora*, as expected from an outcrossing species with commensurately greater effective population size. However, we find that even for *A. thaliana*, which has a low effective recombination rate and for which the effects of linked selection are expected to be strong (Ellegren & Galtier, 2016), local genome features of genome biology have a sufficiently strong impact to affect the shape and scale parameters of the deleterious DFE at the per-gene level, with *A. thaliana* and *C. grandiflora* showing similar patterns of variation.

We took the approach of binning genes by their features and calculating a combined SFS across all genes within each bin in order to have reasonable power to infer the DFE. Our study therefore has some important caveats. We only consider one gene feature at a time in our analyses, and it is possible that co-correlations among variables might drive some of the patterns we observe in our data. While we have attempted to correct for this in our analyses of expression level and gene network connectivity for several commonly studied gene features, it is possible that a feature not included in our analysis has an effect. Approaches that consider the combined effects of gene features are likely to have considerably more power to investigate variation in the DFE and gene-level selective constrain across the genome, however, such approaches make it difficult to disentangle the individual effects of each gene feature. It is also important to note that in analyses of binned data, the number of bins is expected to have an effect, such that the smaller the number of bins, the higher the proportion of variance one expects to explain artefactually, and thus, *R^2^* values should be treated with caution. However, the direction of relationships is still informative, as can be seen by the fact that our relationships between genome biology traits and molecular traits are consistent with different numbers of bins (for a comparison across bins of 10 and 15, see Suppl. Figs. 1 and 2). We also see qualitatively similar patterns across genes with *π_0_*/*π_4_* and genome biology traits.

While we found relationships between genome biology and molecular traits relating to negative selection, our results for positive selection are less clear. Under the FGM and the ‘cost of complexity’ theory, we might expect *α* to decline with complexity (Fisher, 1930; Orr, 2000; Tenaillon, 2014). In *A. thaliana*, the inferred rate of adaptive substitution *α* was very low, and we do not see any relationship with *α* and genome biology traits. This is somewhat in agreement with previous findings; Moutinho et al. (2019) found only a weak relationship in *A. thaliana* between recombination rate and the rate of adaptive substitution, *ω_α_*, but no significant relationship between protein length, expression level or breadth of expression and *ω_α_*. Galtier et al. (2015) also found no relationship between expression level and *α*. It may be that such patterns are generally not observed due to noise, since *α* is difficult to estimate, especially compared to the far more ubiquitous force of purifying selection. It has previously been reported that most adaptive evolution occurs at the surface of proteins, which implies that trends in adaptive evolution are easier to observe in relatively unconstrained protein regions (Moutinho et al., 2019). It also may be that adaptation is focussed on specific sites within genes, which makes it less likely that we will observe general patterns across the genome with *α*. This is supported by Slotte et al. (2011), who found little evidence for adaptive evolution in *A. thaliana* when averaging over the genome, but did find some evidence when considering genes in particular GO categories.

However, in *C. grandiflora*, for which we have evidence of a considerable proportion of adaptive substitutions, we found some indication that genome biology traits related to higher pleiotropy might be positively related to *α*. Although *α* is the proportion of adaptive substitutions, and not the rate of adaptive evolution, this result is still not expected given the ‘cost of complexity’ hypothesis. However, similar findings have been reported in a study of local adaptation in plant species, which found that genes that were more highly connected in gene networks and more broadly expressed tended to be involved in adaptive responses to climate (Whiting et al., 2024). Similarly, genes central to co-expression networks were more likely to undergo adaptive expression level changes following exposure to heat and drought stress in *Tribolium castaneum* beetles (Koch et al., 2025). The same authors also found that eQTLs with higher pleiotropy were under stronger positive selection, which they interpreted as evidence that adaptation may occur via large phenotypic changes produced by changes at highly expressed and highly connected genes, which is somewhat supported by our results.

Our central result, that gene features related to pleiotropy/complexity, including gene network connectivity, number of associated GO terms, and expression level, are positively associated with the strength of deleterious selection acting against new deleterious mutations and, to a lesser extent, negatively associated with the coefficient of variation of selection acting on new mutations, is expected under the FGM for well-adapted populations experiencing stabilising selection at their mutation-selection optima. However, this is not to discount more detailed mechanistic hypotheses about the specific functional effects of mutations and how particular traits affect fitness. As noted by Tenaillon (2014), the great strength of the FGM is its fundamental simplicity and integrative nature, hence its broad applicability. The FGM can be fully described with only three metaparameters: phenotypic complexity, the fitness function, and the way mutations affect the phenotype of an organism. Each of these three metaparameters are complex, such that the FGM integrates over many layers of interactions and molecular mechanisms, with the FGM emerging as a property of more mechanistic and complex models of phenotypic variation (Martin, 2014).

Variation in the DFE has previously been investigated across species, with authors similarly finding some support for the predictions of the FGM (Lin et al., 2025). However, comparing the DFE across species is difficult as species vary hugely in confounding factors, including life history and *N_e_*, the effects of which may be difficult to remove from analyses. Our cross-genome approach reduces the effects of these confounders as the genome generally shares a single taxonomy and has more limited variation in local *N_e_* than exists across species, allowing for a more direct investigation of how parameters of the DFE vary with gene features. We demonstrate here that variation in pleiotropy/complexity among gene modules is sufficient to have demonstrable effects on molecular evolutionary patterns. This analysis can be considered complementary to experimental studies that directly investigate the effects of new mutations, such as the work conducted by Wu et al. (2022), who looked at the impact of mutations, and how these impacts were moderated by expression level, in two genes. Exploring the relationship between genetic architecture and trait evolution at a broad scale remains a promising direction for future research, with recent work incorporating data from GWAS studies to advance our understanding of the relationship between genetic and phenotypic variation (Simons et al., 2025). Such approaches are likely to improve our understanding of how genome features moderate the effects of new mutations in the near future.

## Materials and Methods

### Dataset

This project required species for which there is a well-studied genome and a large amount of polymorphism data, to ensure that we have sufficient power to calculate population genetic summary statistics for relatively small genomic subsets. We therefore opted to conduct our analyses using the 1001 Genomes project dataset of *A. thaliana* (1001 Genomes Consortium, 2016, version 3.1 available at https://1001genomes.org/). We restricted our analyses to the protein-coding portion of the genome. We also used data from the whole genome sequences of 50 *C. grandiflora* individuals. Details for the *C. grandiflora* dataset used in this analysis are available from Mackintosh et al. (2025).

### Ancestral state assignment

For *A. thaliana*, to assign ancestral states to polymorphic sites, we aligned the *A. thaliana* reference genome to two outgroup species, *Arabidopsis lyrata* (genome assembly v.1.0, downloaded from: https://www.ebi.ac.uk/ena/browser/view/GCA_000004255.1) and *Capsella rubella* (genome assembly Caprub1_0, downloaded from: https://www.ebi.ac.uk/ena/browser/view/GCA_000375325.1), using bwa-mem. We then used Est-SFS to assign likely ancestral states to sites, after first randomly down-sampling the SFS to 100 sequenced *A. thaliana* individuals, which is the maximum permitted number for Est-SFS to run, by drawing from a hypergeometric distribution. Zerofold and fourfold synonymous sites were then annotated using our own scripts. For *C. grandiflora*, the genome was aligned with *C. orientalis*, and ancestral states of alleles were estimated by assuming parsimony; any sites that were multi-allelic or at which there was a polymorphism in both species were removed from the dataset. Further details are available from Mackintosh et al. (2025). While ancestral state identification is known to be error prone, we note that our downstream methods include an inference of ancestral error rates, which are reasonable (Suppl. Tables 1 and 2).

### Gene features

For *A. thaliana*, gene features were taken from a variety of sources. Gene network connectivity was estimated as the number of interaction partners each gene had, using the ‘full integrated network’ of AraNet v.2 (Lee et al., 2015, available at https://www.inetbio.org/aranet/), a database of co-functional gene networks. Recombination rates were taken from Brazier and Glémin (2022). For per gene rates, we took rates as estimated from the appropriate 100kb genomic window (available at https://github.com/ThomasBrazier/diversity-determinants-recombination-landscapes-flowering-plants/tree/main/data-cleaned/recombination_maps/loess/100kbwind). Gene length and density were calculated from reference gene and reference genome position information. Gene density was calculated per gene as the average percent CDS in a 100kb genomic window, with the window centred on the midpoint of the gene. Gene expression data was taken from Schmitz et al. (2013, available at https://www.ncbi.nlm.nih.gov/geo/query/acc.cgi?acc=GSE43858; downloaded using the ‘Custom’ option from the GSE43858_RAW.tar archive). Gene expression level was estimated from RNA-seq data and was given in fpkm, fragments per kilobase of exon per million fragments mapped. GO category data for *A. thaliana* was taken from TAIR. We used only those GO classifications that link gene products with biological processes or molecular functions, and discarded GO classifications that link gene products with cellular components or cellular locations.

For *C. grandiflora*, we first identified *A. thaliana* homologs and used the same gene network connectivity estimates and GO term classifications. All other gene features were taken from Josephs et al. (2017). In these experiments, gene expression level was calculated from RNAseq data as the number of read pairs mapping to each gene, normalised by the read count of the entire sample.

### Independent gene features

After finding a number of gene feature traits to be highly correlated, we performed general linear models on two particular gene features of interest: expression level and gene network connectivity. To investigate the effects of each of these gene features, independent of other gene features, we therefore found the best gene feature model to explain the variance in each of our features of interest. We performed model comparisons including a number of gene features such as recombination rate, protein length and gene density to find the simplest model that explained the most variance in our gene feature of interest.

### Best models were as follows

#### A. thaliana

expression level ∼ gene network connectivity + gene density

gene network connectivity ∼ expression level + protein length + interaction of expression level and protein length.

#### C. grandiflora

expression level ∼ gene network connectivity + protein length

gene network connectivity ∼ expression level + protein length

For each of these models, we took the residuals to generate a metric of our gene feature of interest that was independent of other gene features (i.e., for each species, we have an expression level measure that is independent of gene network connectivity, and a gene network connectivity metric that is independent of expression level).

### Population genetics analysis

For our analyses, we first removed splice variants, so that each gene is represented only once. Gene transcripts were chosen based on length, such that the longest transcript was kept, or, if there was no length difference between transcripts, a splice variant was chosen randomly. We subsampled down to 50 haplotypes, to account for missingness in the data and, for *A. thaliana*, we haploidised the genome because we expect there to be an excess of homozygosity in this highly selfing species that will otherwise result in an overrepresentation in even frequency categories of the SFS (Blischak et al., 2020). We then calculated the SFS for zerofold and fourfold synonymous sites per gene. We also calculated *L_0_* and *L_4_*, the total length of zerofold and fourfold sites sequenced per gene, including both monomorphic and polymorphic sites. We then estimated *π_0_* and *π_4_*, zerofold and fourfold synonymous nucleotide diversity, using our own scripts.

We estimated the distribution of fitness effects of new mutations (DFE), using polyDFE (Tataru & Bataillon, 2019). Such an analysis may lack power at the gene level, due to a lack of polymorphisms. Therefore, instead, the DFE was estimates from the SFS as inferred over groups of genes, i.e., different partitions of the genome. For each continuous gene feature, including our independent expression and independent gene network connectivity metric, genes were first ranked by that feature and then divided into groups based on their ranking. Genes were divided such that each group had approximately the same number of 4-fold synonymous polymorphisms (*p_4_*), rather than even numbers of genes. Analyses were performed on genes grouped into 10, 15 and 20 groups, to ensure patterns are robust to different group sizes. For all genome biology features, we then ran polyDFE per group, using the SFS as estimated over all genes in each group, to estimate the full DFE, including the deleterious and beneficial DFE, *α_DFE_*, nuisance parameters and ancestral error.

All statistical analyses and plotting were conducted in R.

## Acknowledgements

This work was supported by the SciLifeLab and Wallenberg Data Driven Life Science Program (grant: KAW 2024.0159), and by a SCAS Natural Sciences Fellowship. The authors would also like to thank Alex Mackintosh for providing the *Capsella* polymorphism data used in this work.

## Data Availability

All python and R code used in this analysis, and intermediate files, are available at https://github.com/j-e-james/GenomeOrganisationSelectionEfficiency.

