## Supplementary material for "The effect of genome organisation on selection efficiency in two contrasted plant species": Suppl. Fig.

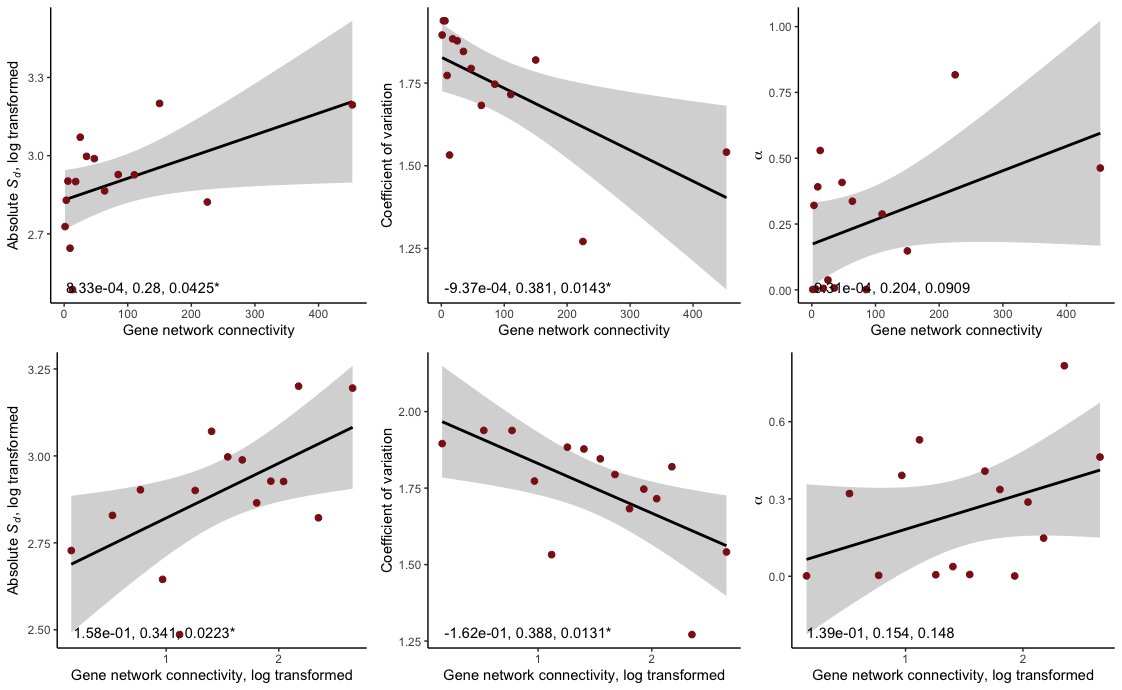

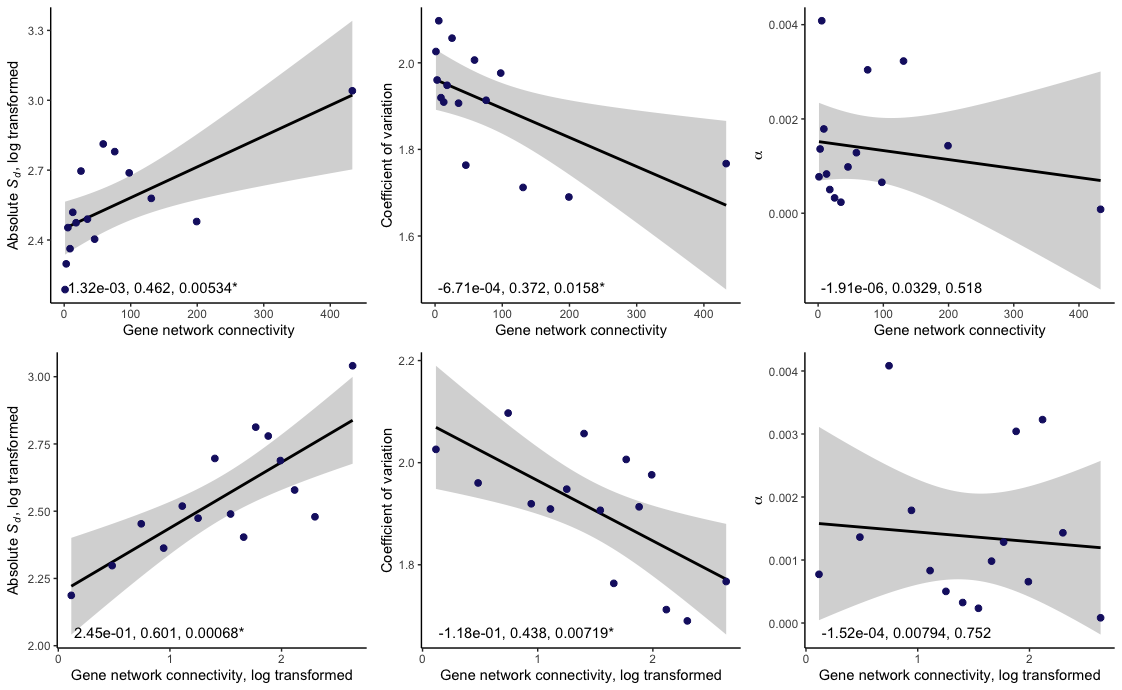

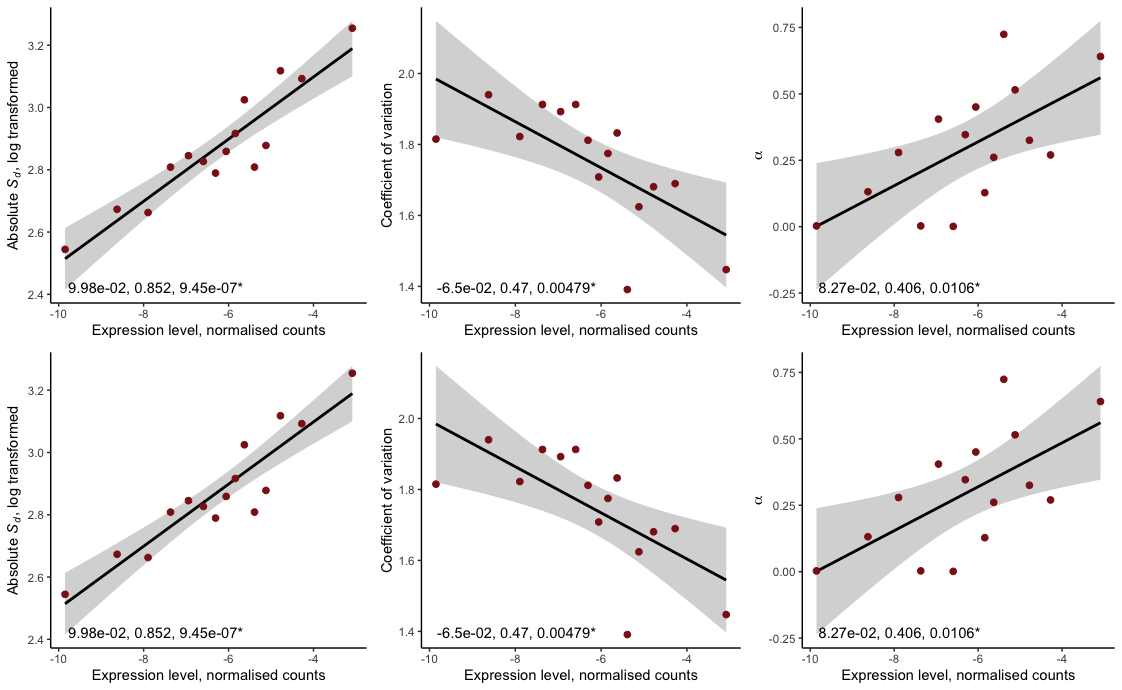

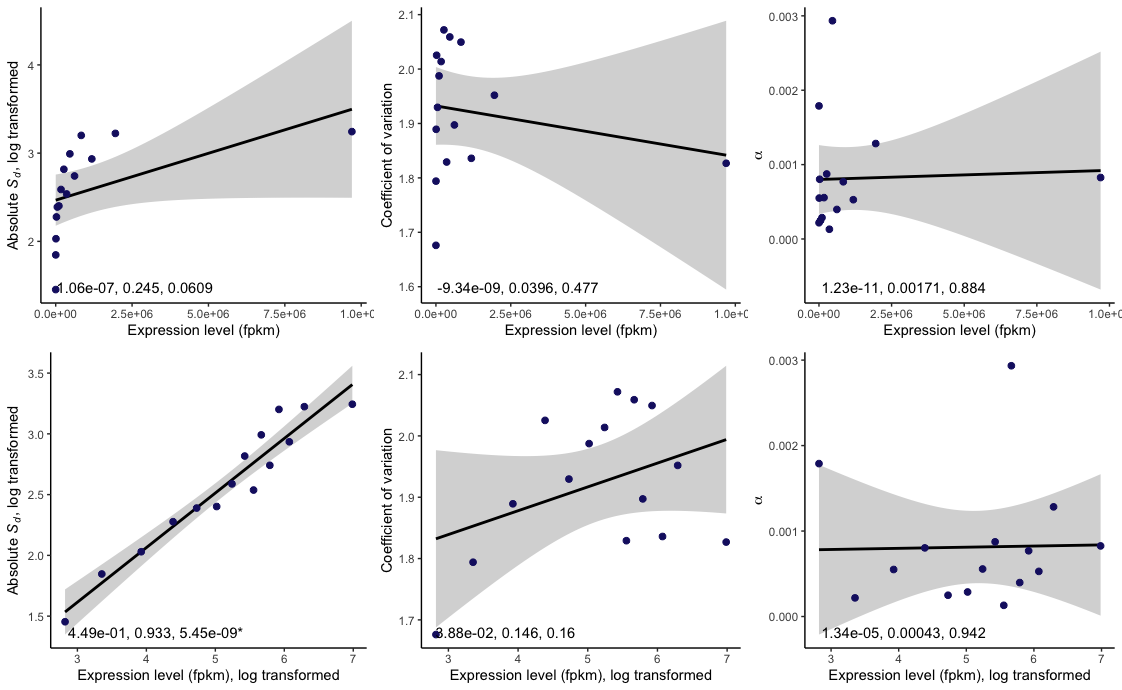

Suppl. Fig. 1) Parameters of the DFE and two genome traits related to pleiotropy; expression level and gene network connectivity. Details are as for main figure 1, shown for 15 bins (main text: 20 bins).The first column shows the absolute mean strength of selection acting against new deleterious mutations, *S_d_* (the scale parameter of the DFE), the second shows the coefficient of variation of selection acting against new deleterious mutations (related to the shape parameter *b* of the DFE), the third shows *α*, the proportion of substitutions estimated to be fixed by adaptive evolution, as estimated from the beneficial DFE (i.e., using polymorphism data only). Rows indicate different genome traits: either expression level or the estimated level of connectedness in functional gene interaction networks, estimated from number of gene interaction partners. Colour indicates species; dark blue: *A. thaliana*, dark red: *C. grandiflora*. Points show the estimated value of the estimated DFE parameter, estimated over all genes in that bin, plotted against the mean value of the genome trait for that bin. Lines are linear regression results (log transformed genome biology traits), shown with 95% confidence intervals. Results of linear regression analyses are shown on each plot (slope, *R^2^*, and *p* value), with results significant at the p < 0.05 level marked ‘*’.

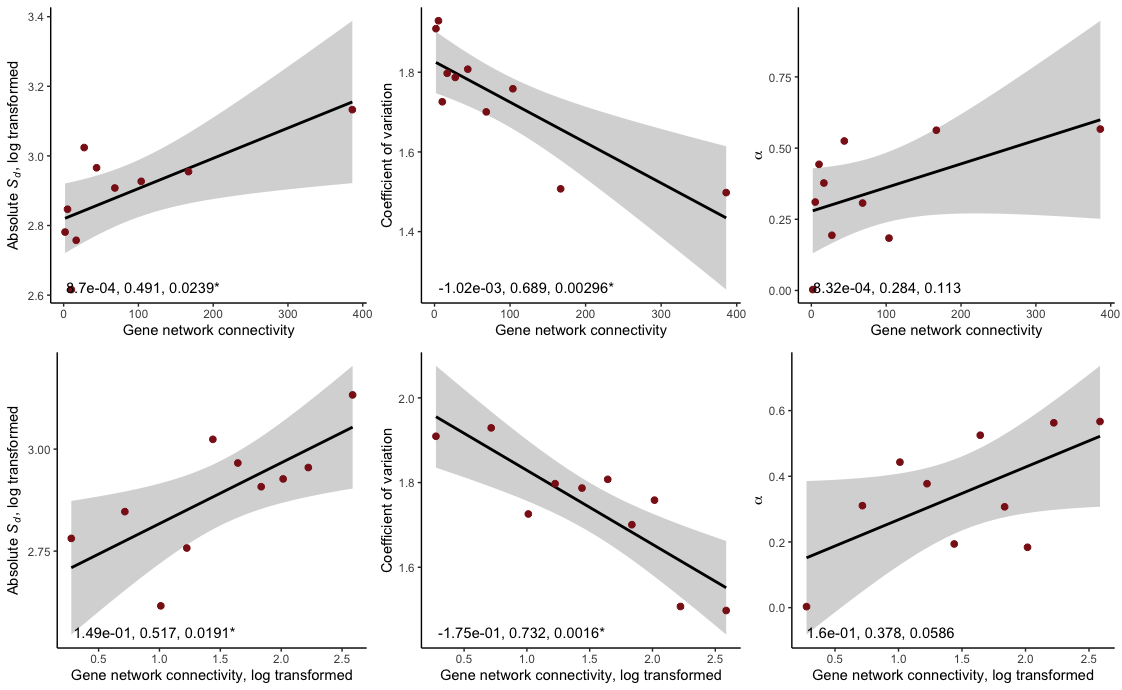

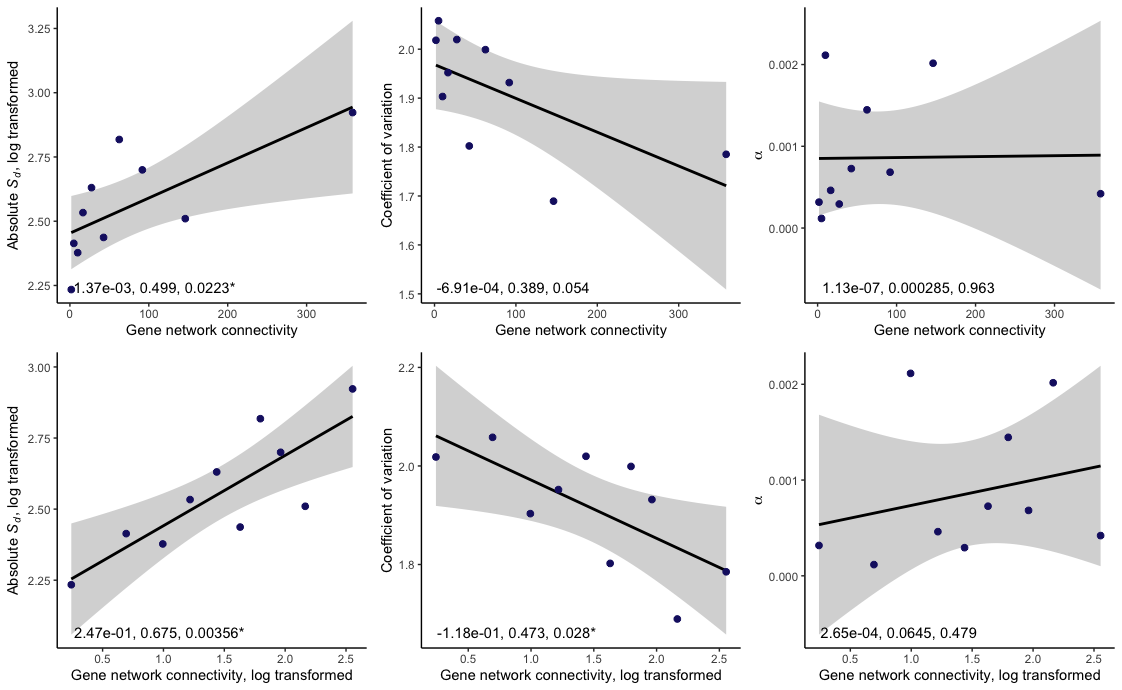

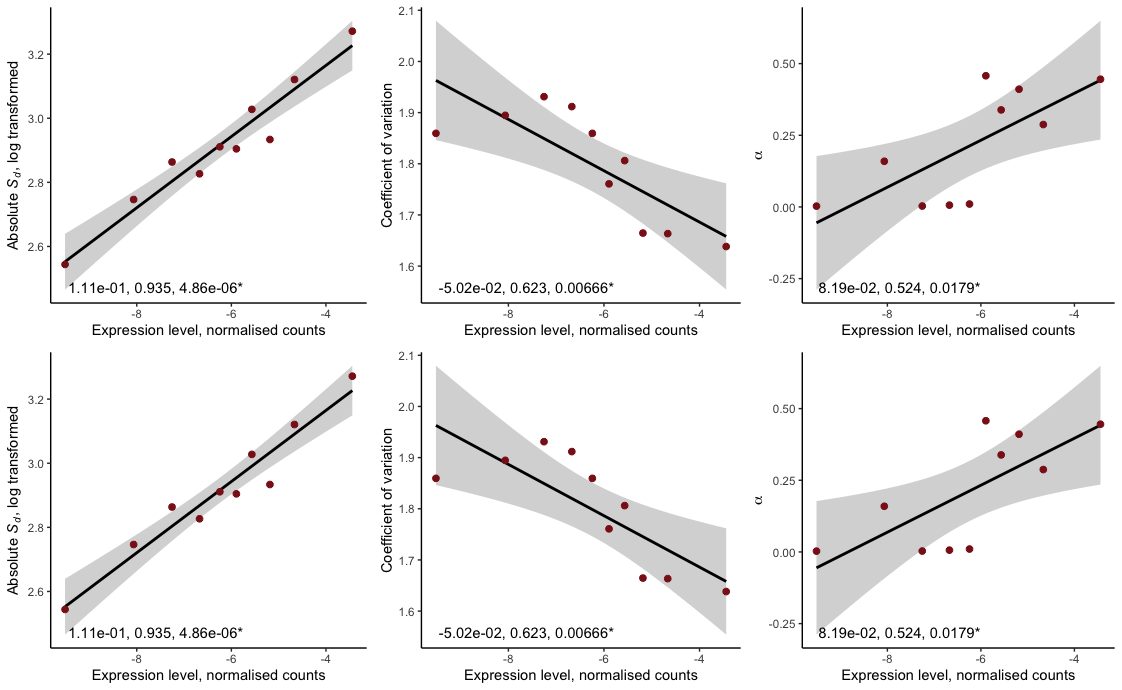

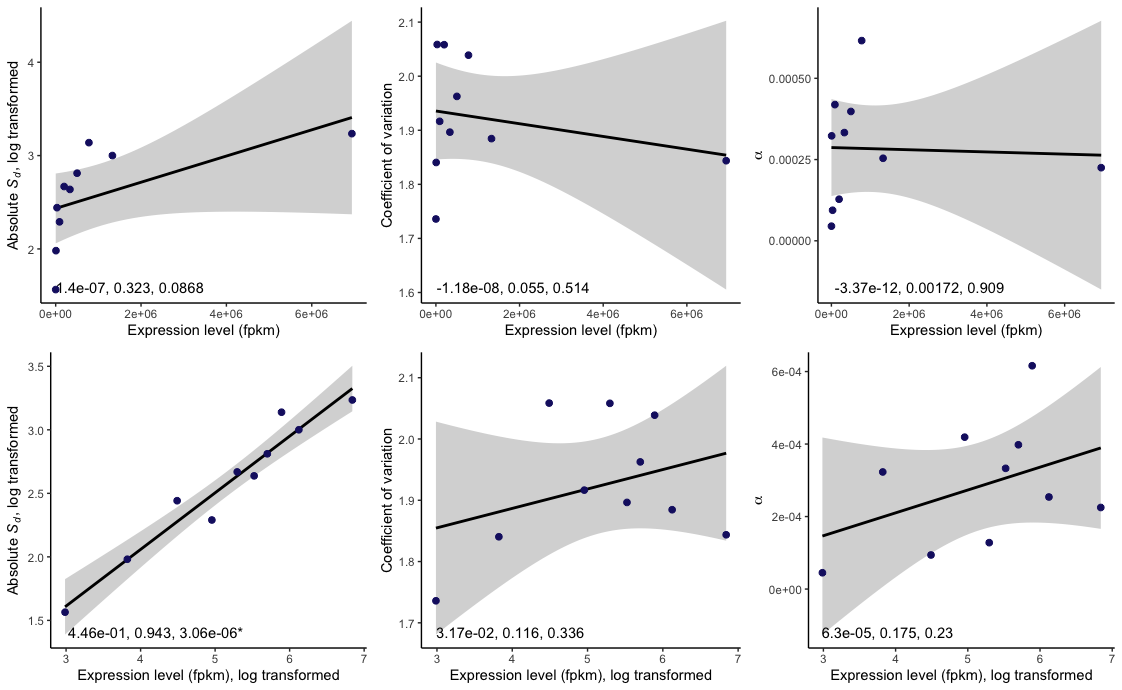

Suppl. Fig. 2) Parameters of the DFE and two genome traits related to pleiotropy; expression level and gene network connectivity. Details are as for main figure 1, shown for 150 bins (main text: 20 bins). The first column shows the absolute mean strength of selection acting against new deleterious mutations, *S_d_* (the scale parameter of the DFE), the second shows the coefficient of variation of selection acting against new deleterious mutations (related to the shape parameter *b* of the DFE), the third shows *α*, the proportion of substitutions estimated to be fixed by adaptive evolution, as estimated from the beneficial DFE (i.e., using polymorphism data only). Rows indicate different genome traits: either expression level or the estimated level of connectedness in functional gene interaction networks, estimated from number of gene interaction partners. Colour indicates species; dark blue: *A. thaliana*, dark red: *C. grandiflora*. Points show the estimated value of the estimated DFE parameter, estimated over all genes in that bin, plotted against the mean value of the genome trait for that bin. Lines are linear regression results (log transformed genome biology traits), shown with 95% confidence intervals. Results of linear regression analyses are shown on each plot (slope, *R^2^*, and *p* value), with results significant at the p < 0.05 level marked ‘*’.

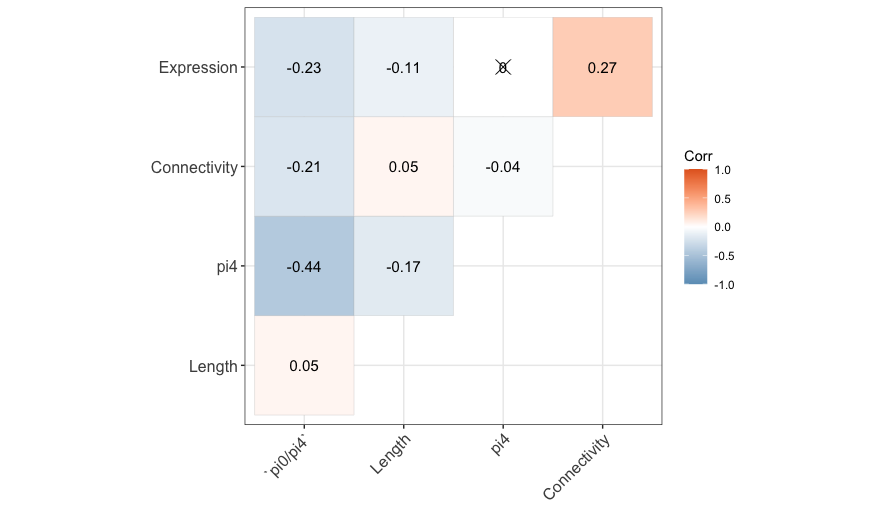

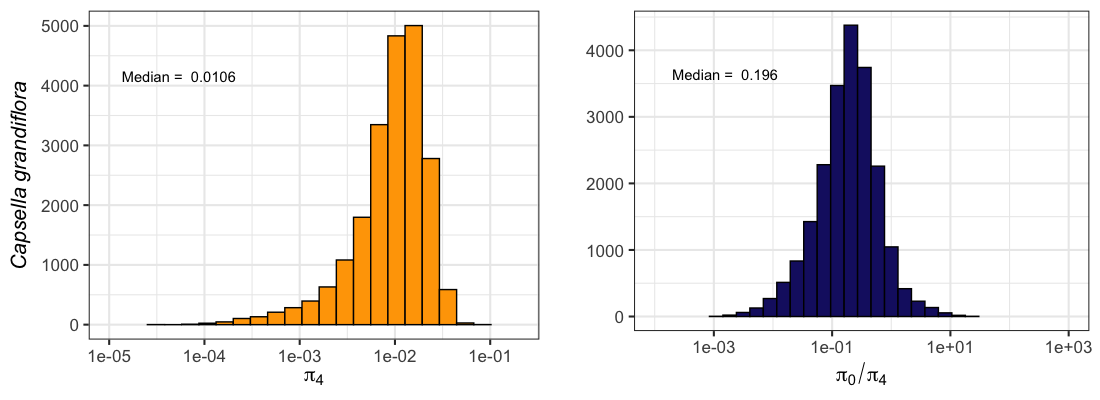

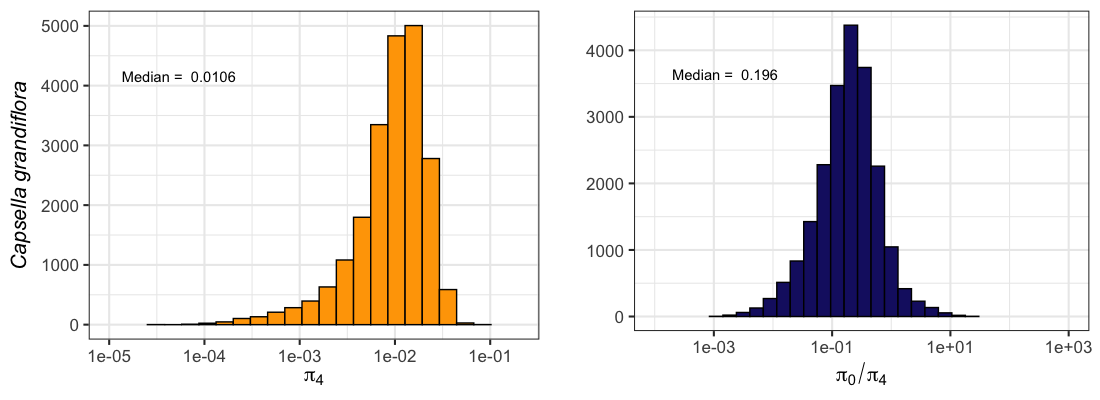

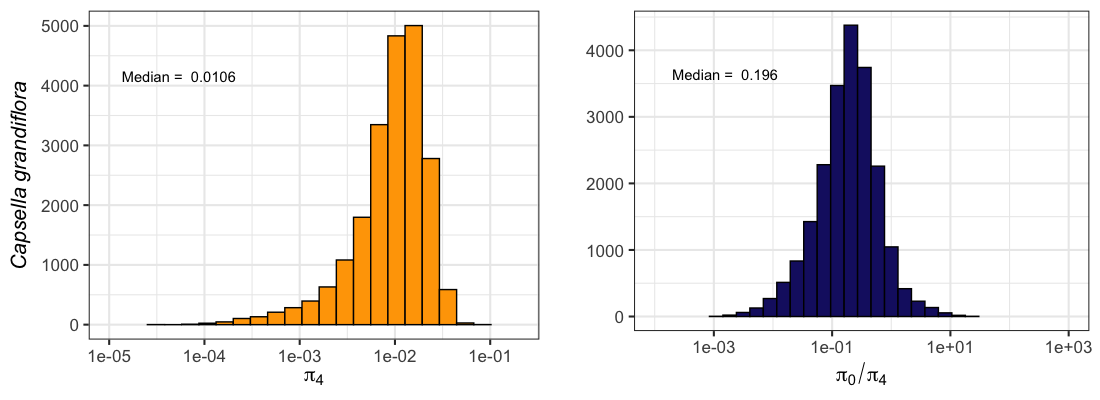

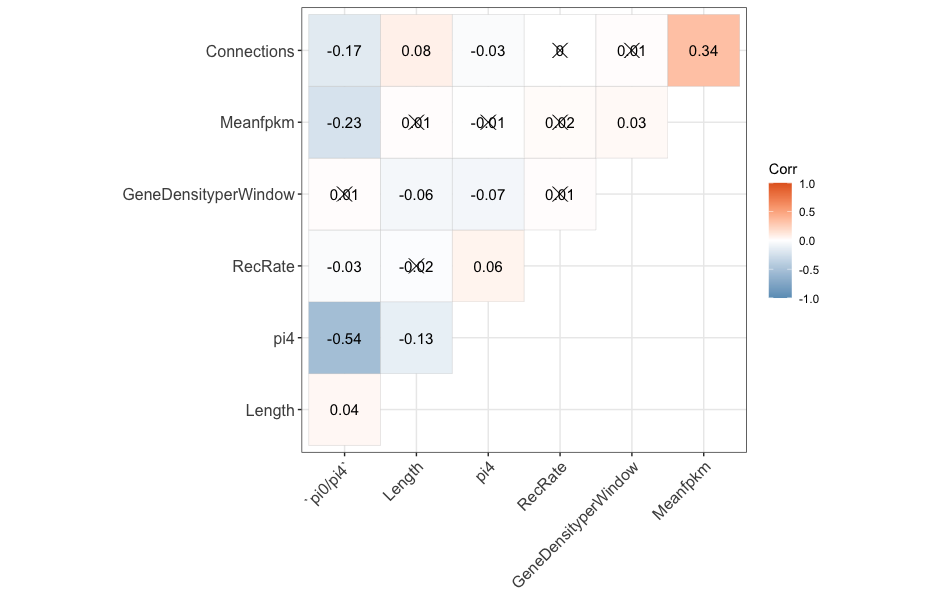

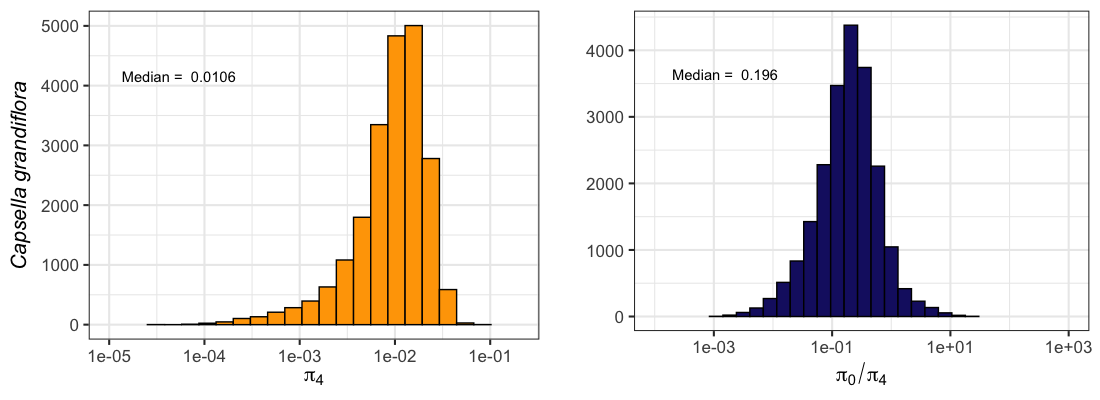

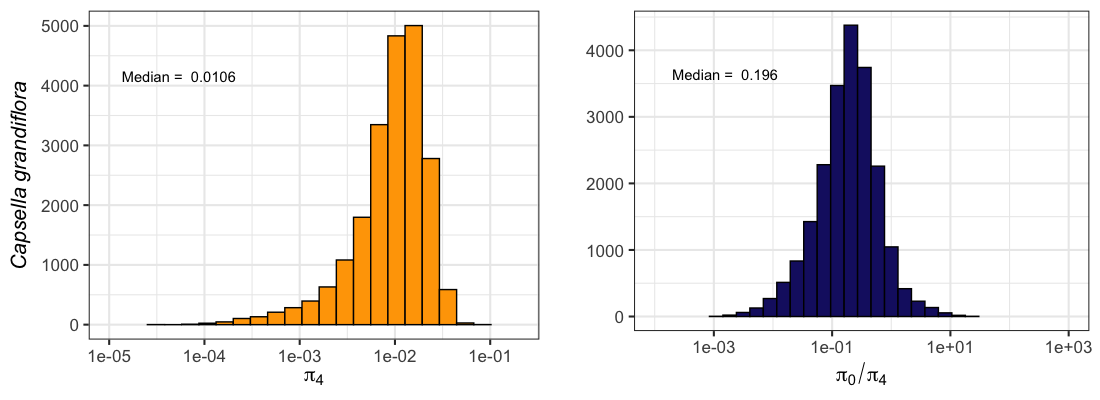

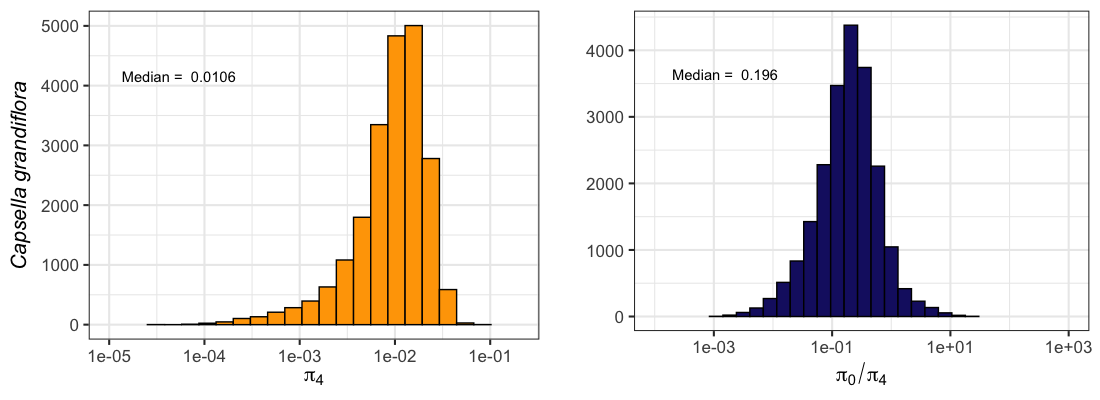

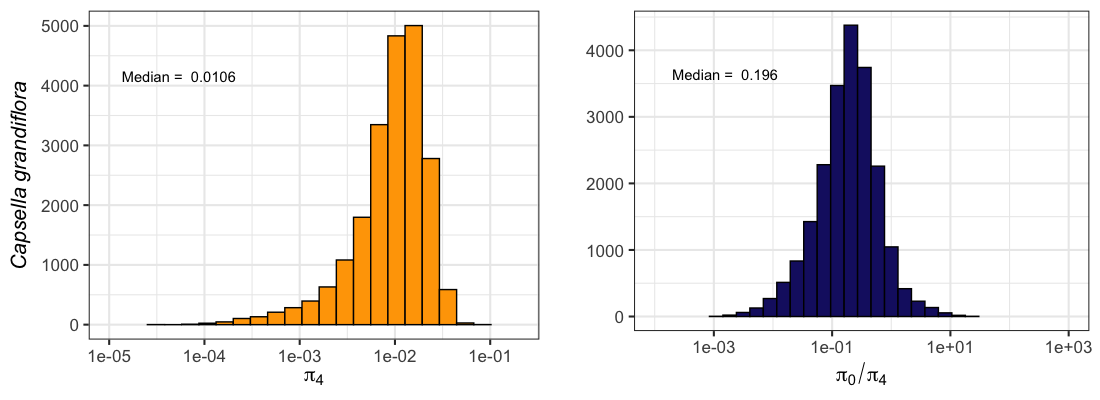

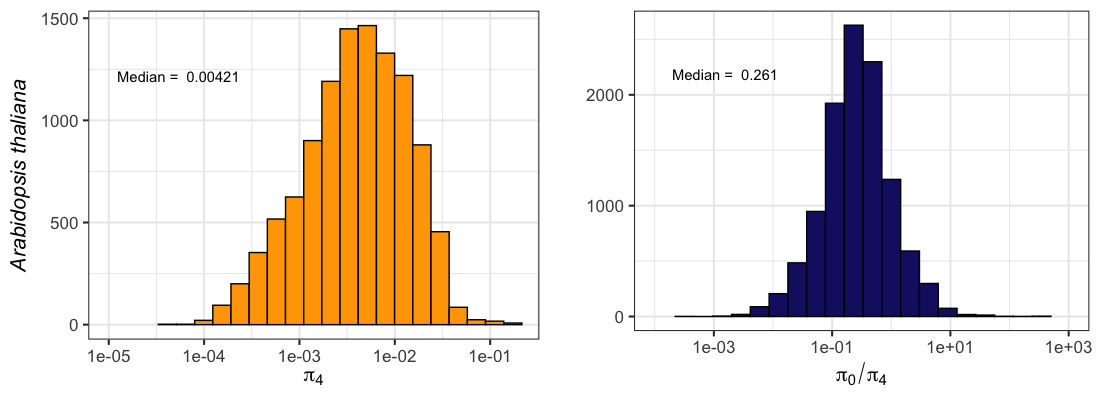

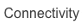

Suppl. Fig. 3) Pairwise correlation matrix of gene-level estimates of genome biology traits, π_4_ and π_0_/π_4_ across the *A. thaliana* and *C. grandiflora* genomes. For *A. thaliana*, all features were log transformed, for C. grandiflora, all features were log transformed except for expression level. Genome biology traits are shortened for ease of viewing: these correspond to: Connectivity: Gene network connectivity, Meanfpkm: Expression level (fpkm), GeneDensityPerWindow: Density of genes in 10kb windows, centred on the gene of interest, RecRate: Recombination rate, Expression; Expression level, normalised counts.
